# A yeast modular cloning (MoClo) toolkit expansion for optimization of heterologous protein secretion and surface display in *Saccharomyces cerevisiae*

**DOI:** 10.1101/2023.12.12.570949

**Authors:** Nicola M. O’Riordan, Vanja Jurić, Sarah K. O’Neill, Aoife P. Roche, Paul W. Young

## Abstract

*Saccharomyces cerevisiae* is an attractive host for expression of secreted proteins in a biotechnology context. Unfortunately, many heterologous proteins fail to enter, or efficiently progress through, the secretory pathway, resulting in poor yields. Similarly, yeast surface display has become a widely used technique in protein engineering but achieving sufficient levels of surface expression of recombinant proteins is often challenging. Signal peptides (SPs) and translational fusion partners (TFPs) can be used to direct heterologous proteins through the yeast secretory pathway, however, selection of the optimal secretion promoting sequence is largely a process of trial and error. The yeast modular cloning (MoClo) toolkit utilises Type IIS restriction enzymes to facilitate efficient assembly of expression vectors from standardized parts. We have expanded this toolkit to enable the efficient incorporation of a panel of sixteen well-characterized SPs and TFPs and five surface display anchor proteins into *S. cerevisiae* expression cassettes. The secretion promoting signals were validated using five different proteins of interest. Comparison of intracellular and secreted protein levels revealed the optimal secretion promoting sequence for each individual protein. Large, protein of interest-specific variations in secretion efficiency were observed. SP sequences were also used with the five surface display anchors and the combination of SP and anchor protein proved critical for efficient surface display. These observations highlight the value of the described panel of MoClo compatible parts to allow facile screening of SPs, TFPs and anchor proteins for optimal secretion and/or surface display of a given protein of interest in *S. cerevisiae*.

**GRAPHICAL ABSTRACT:** 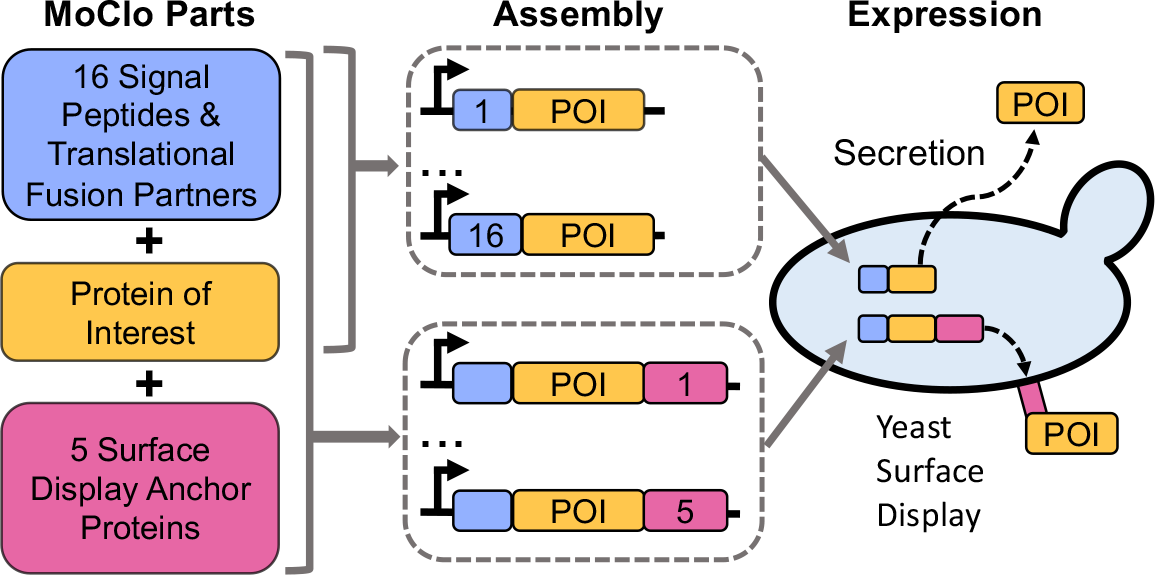

## INTRODUCTION

*Saccharomyces cerevisiae* has been used extensively as a host organism for heterologous protein expression due to factors such as amenability to large scale fermentation, capacity for eukaryotic post-translational modifications and well characterised genetics. Its “generally recognized as safe” (GRAS) status is especially important for proteins with applications in the food and biopharmaceutical sectors. Lower costs and decreased risk of viral contamination are potential advantages of *S. cerevisiae* over mammalian cells for therapeutic protein and antibody production, for example *(1, 2)*. In addition, *S. cerevisiae* based yeast display technology has found many research and biotechnological applications for protein engineering *(3)*. Nevertheless, many proteins are difficult to express at high levels in *S. cerevisiae* and achieving efficient secretion and/or cell surface display of heterologous proteins is often challenging *(3, 4)*.

Protein secretion in *S. cerevisiae* is usually initiated by an amino terminal signal peptide (SP) that is ultimately cleaved from the mature protein. SPs direct secreted proteins into the endoplasmic reticulum (ER) via either post-translational or co-translational translocation mechanisms (e.g. the Sec- and SRP-dependent pathways respectively). The pre-region of a SP precedes the signal peptidase cleavage site and has a tripartite structure generally consisting of a positively charged N-region, an hydrophobic α-helical H-region and the C-region containing the cleavage site (often after an Ala residue) *(5)*. For many proteins a pro-region, after the signal peptidase cleavage site, is also important for efficient secretion. The pro-region is typically removed from the mature protein by additional proteases, such as the Golgi-resident KEX2 protease, for example *(5, 6)*.

Significant efforts have been made to improve the secretion of recombinant proteins by *S. cerevisiae* including various strategies to identify sequences that would direct efficient secretion of a wide range of recombinant proteins. Mori *et al*. (2015) *(7)* screened a library of 60 *S. cerevisiae* SPs and identified six that promoted high levels of secretion of an exogenous β-galactosidase enzyme. Holec *et al*. (2022) *(8)* conducted a proteome-scale screen to identify SPs that gave consistent, high-level surface display of proteins with diverse amino-terminal sequences. Other studies have optimized the widely used α-factor mating pheromone pre-pro-SP through a variety of strategies to achieve enhanced secretion of diverse proteins of interest *(9-12)*. Bae and co-workers *(13)* conducted a genome-wide screen to identify sequences termed translational fusion partners (TFPs) that enhanced the expression and/or secretion of several difficult to produce proteins. The TFPs identified typically contain additional amino acid sequences beyond the amino terminal SP that may promote the correct folding of the recombinant protein to which they are fused or otherwise promote its progression through the secretory pathway. A variety of strategies have also been employed to improve cell surface display of proteins of interest. These include signal peptide optimization and varying the choice of display anchor protein used (reviewed in *(3)*). Overall, while the aforementioned studies identify many promising approaches that can greatly enhance secretion or cell surface display in *S. cerevisiae*, no one strategy is likely to be optimal for every heterologous protein of interest.

In the absence of a universal secretion promoting sequence that will work for all recombinant proteins, an efficient system to rapidly sample a library of such sequences is highly desirable. The same principle applies to the selection of anchor proteins for yeast display. Modular cloning (MoClo) is a standardized, hierarchical system that facilitates the efficient assembly of plasmids containing single or multiple transcriptional units using type IIS restriction enzymes *(14)*. The MoClo strategy has been widely adopted *(15)* and has also been adapted for specific applications including for use in *S. cerevisiae* with the development of a modular cloning yeast toolkit (MoClo YTK) *(16)*. Here we have formatted a variety of secretion enhancing and surface display sequences described in the scientific literature for use in conjunction with the yeast MoClo toolkit in a way that permits the production of recombinant proteins with native (or near native) amino terminal sequences following SP or TFP cleavage. The resulting collection of plasmids constitutes a yeast secretion/display (YSD) toolkit that will be made available to the research community.

## RESULTS AND DISCUSSION

### Expansion of the MoClo YTK to accommodate a panel of secretion promoting sequences

The MoClo yeast toolkit includes a selection of promoter and terminator elements that can be combined with a coding sequence to generate transcriptional units for protein expression in vectors with several possible selectable markers and origins of replication. We sought to expand this toolkit by developing SP and TFP sequences as “parts” that can be readily combined with a coding sequence of interest to optimize protein secretion or surface display (Fig 1A). Maintaining the native amino- and/or carboxyl-terminal amino acid sequence can be critical to the functionality of recombinantly expressed proteins and may also be important for regulatory reasons. With this in mind we considered the sequence of yeast SP pre-sequences, which most frequently have an alanine residue at the -1 position relative to the signal peptidase cleavage site as part of an Ala-X-Ala or Val-X-Ala motif *(17)*. To join SP pre-sequences ending with Ala to coding sequences, without introducing any extra amino acids, we chose an TGCT overhang for DNA assembly in which the GCT codes for Ala (Fig 1B). A TGCT overhang is predicted to have high fidelity when used as part of the YTK toolkit when analysed using the NEBridge GetSet tool (Supp Fig 1). This approach ensures that coding sequences with their native amino termini will be generated following signal peptidase cleavage of the SP pre-sequence.

**Figure 1.**
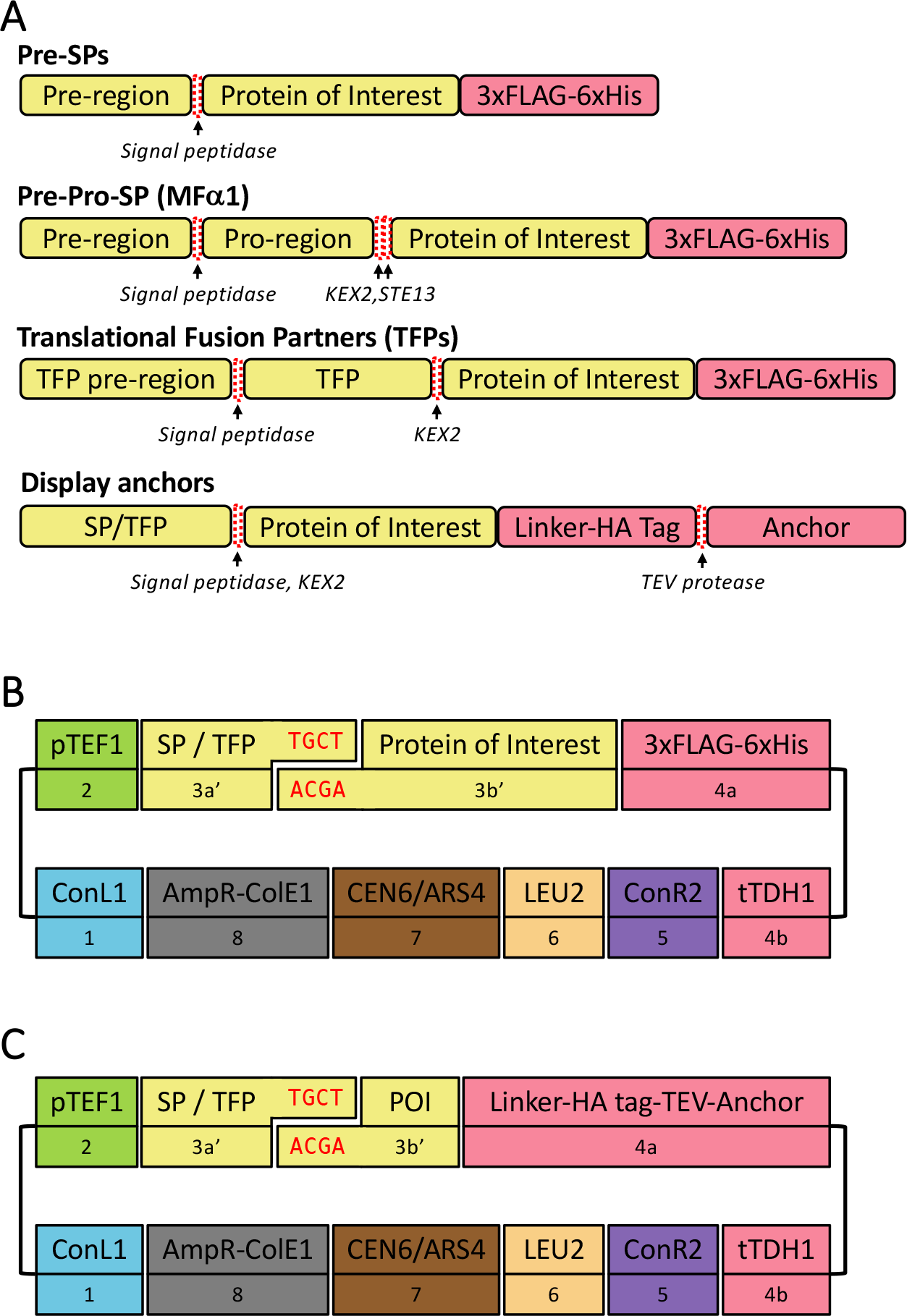
Schematic representation of constructs and assembly strategy. **(A)** The indicated types of amino-terminal secretion promoting sequences were combined with a protein of interest and a carboxyl-terminal 3xFLAG-6xHistidine tag to assess their secretion efficiency, or a HA epitope tagged anchor protein for surface display experiments. **(B)** Yeast expression constructs to evaluate a panel of SPs and TFPs were assembled using the standard parts from the yeast modular cloning toolkit with the exception of the parts denoted as 3a’ and 3b’ that are joined by a custom overhanging sequence (red text). **(C)** Similar expression constructs were used to evaluate yeast surface display anchors which were assembled as standard 4a part types and encoded a flexible linker (GGGGSGGGGS), an HA epitope tag and a TEV protease cleavage site upstream of the anchor protein. POI = Protein of interest

We initially selected a panel of nine SP pre-sequences, three TFPs and an engineered mating factor α pre-pro-sequence (MFαpp8) that was previously reported to enhance antibody secretion *(11)* (Table 1). The selected sequences were designed as yeast modular cloning parts that are equivalent to subtype 3a parts according to the convention of Lee at al *(16)* except using a TGCT rather than TTCT at the 3a/3b junction (Fig 1B). We denote these parts as 3a’ and the coding sequence parts as 3b’ to highlight this distinction. The choice of TGCT overhang necessitated changing glycine to alanine at the -1 position for two SP pre-sequences and an amino acid change at the -2 position for one of these (Table 1). For the MFαpp8 sequence an Glu-Ala to Asp-Ala change was made for similar reasons at the second STE13 protease cleavage site. A recognition sequence for KEX2 protease (Leu-Asp-Lys-Arg) was included after the TFP sequences such that a recombinant protein with just an additional amino-terminal alanine is expected to be generated following proteolytic cleavage.

**Table 1.**
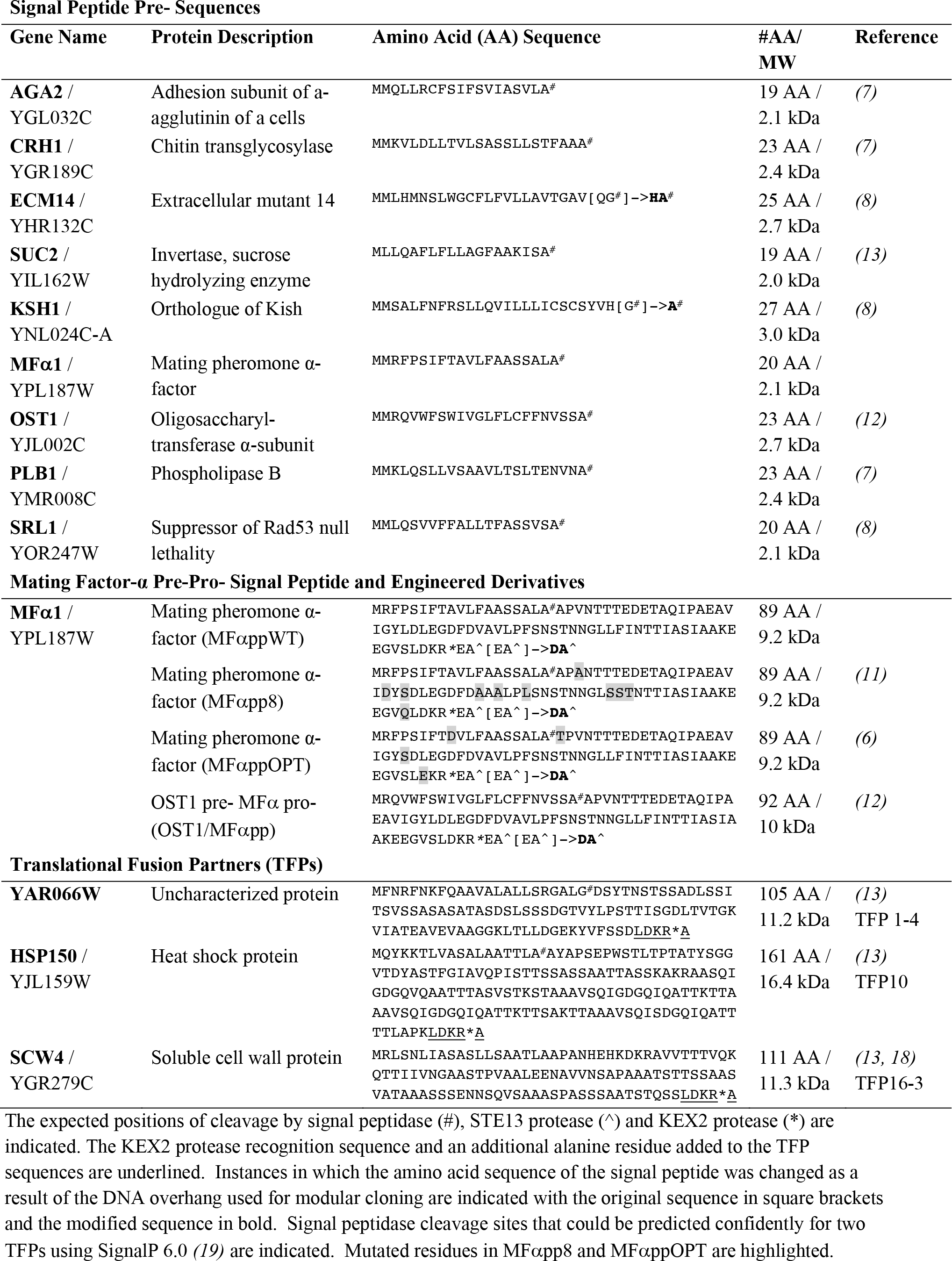
Signal Peptides and Translational Fusion Partners used in this study.

### Evaluation of the panel of secretion promoting sequences

Expression vectors were assembled for each SP and TFP using a strong constitutive pTEF1 promoter (Fig 1B). A type 4a part encoding a carboxyl-terminal 3xFLAG-6xHistidine tag was included to facilitate detection and/or purification of the recombinant protein. As an initial test of the system the expression of a known *S. cerevisiae* secreted protein, invertase (SUC2), was evaluated by western blotting of culture media. Secreted invertase was detected for all SP and TFP sequences at a very high molecular weight (Supp Fig 2). This is likely due to extensive glycosylation as previously reported *(20, 21)*). This indicated that all the SPs and TFPs were functional when used in conjunction with a readily secreted protein.

We then tested the system for production of nanobodies – variable domains of heavy-chain-only antibodies from camelid species that have widespread applications as imaging reagents and therapeutics. An anti-green fluorescent protein (GFP) nanobody *(22)* was readily secreted into the culture media using all the SPs and TFPs and was detected as a broad band by western blotting – possibly a consequence of differing levels of glycosylation (Fig 2A). Similarly, differential glycosylation, or altered cleavage at protease recognition sites, may explain the slightly increased molecular weight of the secreted nanobody observed with the MFαpp8 SP and the YAR066W TFP. Intracellularly, for all pre-SPs the nanobody was detected as a relatively discrete band at the expected ∼18 kDa molecular weight of the pre-protein with a less prominent band of slightly lower molecular weight potentially representing the nanobody following pre-SP removal by signal peptidase cleavage. The MFαpp8 SP, and three TFPs resulted in multiple forms of the intracellular nanobody resolving at higher molecular weights. These forms likely represent proteins in which the pre-pro-SP or TFP is uncleaved or partially cleaved and have additionally been glycosylated. Bands of molecular weight less than 18 kDa probably indicate some intracellular degradation of nanobody. When purified from the media the nanobody was able to efficiently bind to GFP in a pull down assay – verifying its functionality (Fig 2A, lower panel).

**Figure 2.**
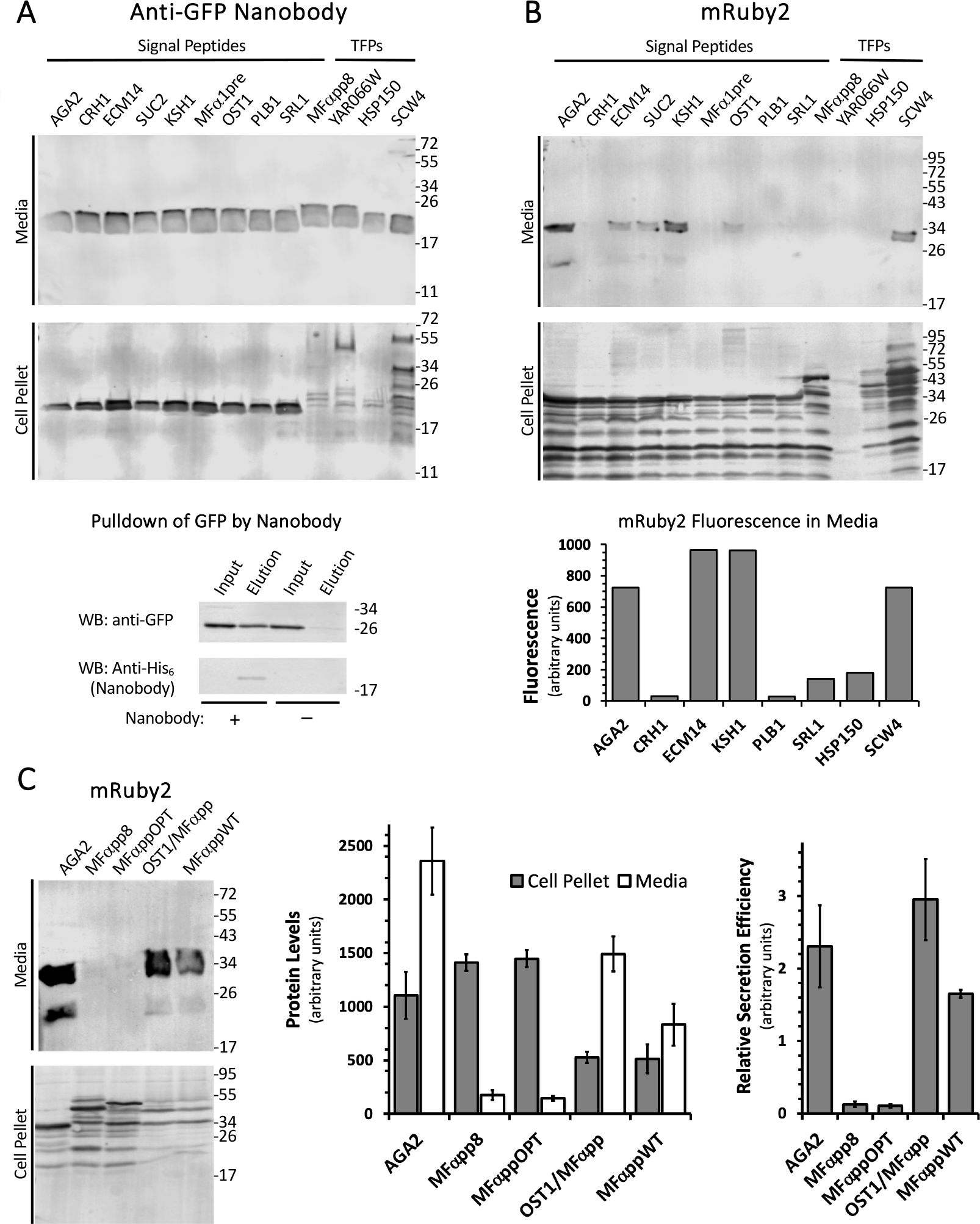
Evaluation of a panel of secretion promoting sequences for the expression of a nanobody and the mRuby2 fluorescent protein. Recombinant proteins were targeted for secretion using the indicated SPs and TFPs. Secreted and intracellular proteins were detected from the media and cell pellet respectively by western blotting for a carboxyl-terminal 6xHis tag. **(A)** An anti-GFP nanobody was readily secreted using all SPs and TFPs (upper panel) and was functional as assessed by its ability to pull down GFP from a cell lysate of GFP-expressing HEK293T cells (lower panel). (**B**) Only half of the SPs and one TFP directed efficient secretion of the mRuby2 fluorescent protein as assessed by western blotting (upper panel). Fluorescence measurements of cell-free media confirmed this pattern for a subset of the SPs and TFPs (lower panel). **(C)** Wildtype and engineered variants of the MFα pre-pro-SP differ significantly in their ability to direct secretion of mRuby2 as assessed by western blotting (left panel). Protein levels from three independent experiments were quantified (middle panel) and the ratio of levels observed in media versus cell pellet samples was calculated as a measure of relative secretion efficiency (right panel). Sizes of molecular weight markers are indicated in kDa. Error bars represent SEM.

Next we examined the secretion of the fluorescent protein mRuby2 *(23)* (Fig 2B). In contrast to the nanobody, efficient secretion of mRuby2 was only observed with the pre-SPs AGA2, ECM14, SUC2, KSH1 and OST1 and the TFP SCW4. Intracellularly, relatively uniform levels of mRuby2 was detected for all SPs and TFPs except YAR066W, albeit that multiple lower molecular weight bands suggestive of protein degradation are observed. Differences in mRuby2 levels in the media therefore seem to reflect the inability of some pre-SPs and TFPs to direct mRuby2 through the secretory pathway rather than a failure of protein expression. Secreted mRuby2 was observed as a doublet band at the correct molecular weight of ∼33 kDa. Whether the two bands result from partial proteolysis or post-translational modification is not clear. However, the secreted protein is functional as indicated by fluorescence measurements of the media in separate experiments using a subset of the SPs and TFPs, with the same pattern of secretion efficiency being apparent (Fig 2B, lower panel).

Neither the MFα1 pre-nor the engineered MFα1 pre-pro-sequence MFαpp8 directed efficient secretion of mRuby2. Several other improved MFα1 pre-pro-sequences have been described in the literature. We were curious to examine the performance of two of these in promoting secretion of mRuby2 in comparison to MFαpp8 and the wildtype MFα1 pre-pro-sequence (Table 1). MFαppOPT arose from a systematic attempt to design of a universal signal peptide *(6)* and has four amino acid substitutions compared to the wildtype sequence. By contrast, OST1-MFαpp is a fusion of the OST1 pre-sequence with the MFα1 pro-sequence that was used to promote secretion of a monomeric superfolding GFP variant *(12)* and three distinct exogenous proteins in *S. cerevisiae (9)* via co-translational targeting to the ER. Secretion of mRuby2 directed by the wildtype MFα pre-pro-SP, MFαppOPT, MFαpp8 and OST1/MFαpp was compared to the AGA2 pre-SP – the most efficient sequence from our initial panel. OST1/MFαpp and MFαppWT directed reasonable levels of mRuby2 secretion into the media, though not as high as for AGA2. Only minimal secretion was observed for MFαpp8 and MFαppOPT. Protein levels in the cell pellet were lower for OST1/MFαpp and MFαppWT compared to the other sequences. Calculating the ratio of mRuby2 levels in the media versus the cell pellet as a measure of relative secretion efficiency revealed OST1/MFαpp to be on a par with AGA2, while MFαppWT was slightly less efficient but still dramatically better than either of the engineered MFαpp8 and MFαppOPT sequences.

The profiles of secretion observed for the anti-GFP nanobody and mRuby2 differ significantly. The former was efficiently secreted regardless of the secretion promoting sequence employed. By contrast, only six of thirteen sequences in our initial panel directed appreciable secretion of the mRuby2 fluorescent protein with the pre-SPs AGA2, and KSH1 and TFP, SCW4 giving the highest levels of secretion. Secretion by *S. cerevisiae* of mRuby2 has not, to our knowledge, been previously described in the literature. However, difficulties in achieving secretion of GFP from *S. cerevisiae* have been reported *(24)* and it has been suggested that SPs, such as OST1, that direct the recombinant protein towards the co-translational translocation pathway at the ER are preferable for secretion of some fluorescent proteins *(12)*. While the OST1 pre-SP directed low levels of secretion of mRuby2 in this study, the hybrid OST1/MFαpp was more successful. Surprisingly, the engineered MFαpp8 and MFαppOPT sequences, that have been shown to enhance secretion of numerous proteins, failed to direct efficient secretion of mRuby2, while the wildtype sequence from which they were derived was relatively successful. MFαpp8 in particular, has been routinely used in place of the wildtype sequence, but our observations caution against the assumption that engineered SPs are always an improvement over wildtype sequences for every protein of interest.

### Application of the toolkit expansion for secretion of sweet proteins

Sweet proteins are potential low calorie alternatives to sugar and artificial sweeteners and *S. cerevisiae* is and attractive host for their production given its widespread use in the food and beverage industry *(25)*. We therefore applied the panel of pre-SPs and TFPs for the secretion of two sweet proteins brazzein, *(26)* and monellin *(27)*. Brazzein was expressed at rather low levels overall, nonetheless, all SPs and TFPs from the panel except for HSP150 promoted brazzein secretion (Fig 3A). The secreted protein was observed as a broad band running at, or just above, the 17 kDa molecular weight marker higher – somewhat higher than its predicted ∼10 kDa molecular weight. Intracellularly, when expressed with the pre-SPs, brazzein was generally detected as either a single well-defined band or as a doublet (for ECM14 and KSH1), just below the 17 kDa marker. Taken together, this suggests that the broad band observed for secreted brazzein may reflect differentially glycosylated states of the protein. Little to no intracellular protein was detected for HSP150, explaining the absence of secreted brazzein for this TFP. Interestingly, high levels of intracellular protein for YAR066W compared to the SCW4 TFP or the pre-SPs, for example, did not result in greater brazzein secretion.

**Figure 3.**
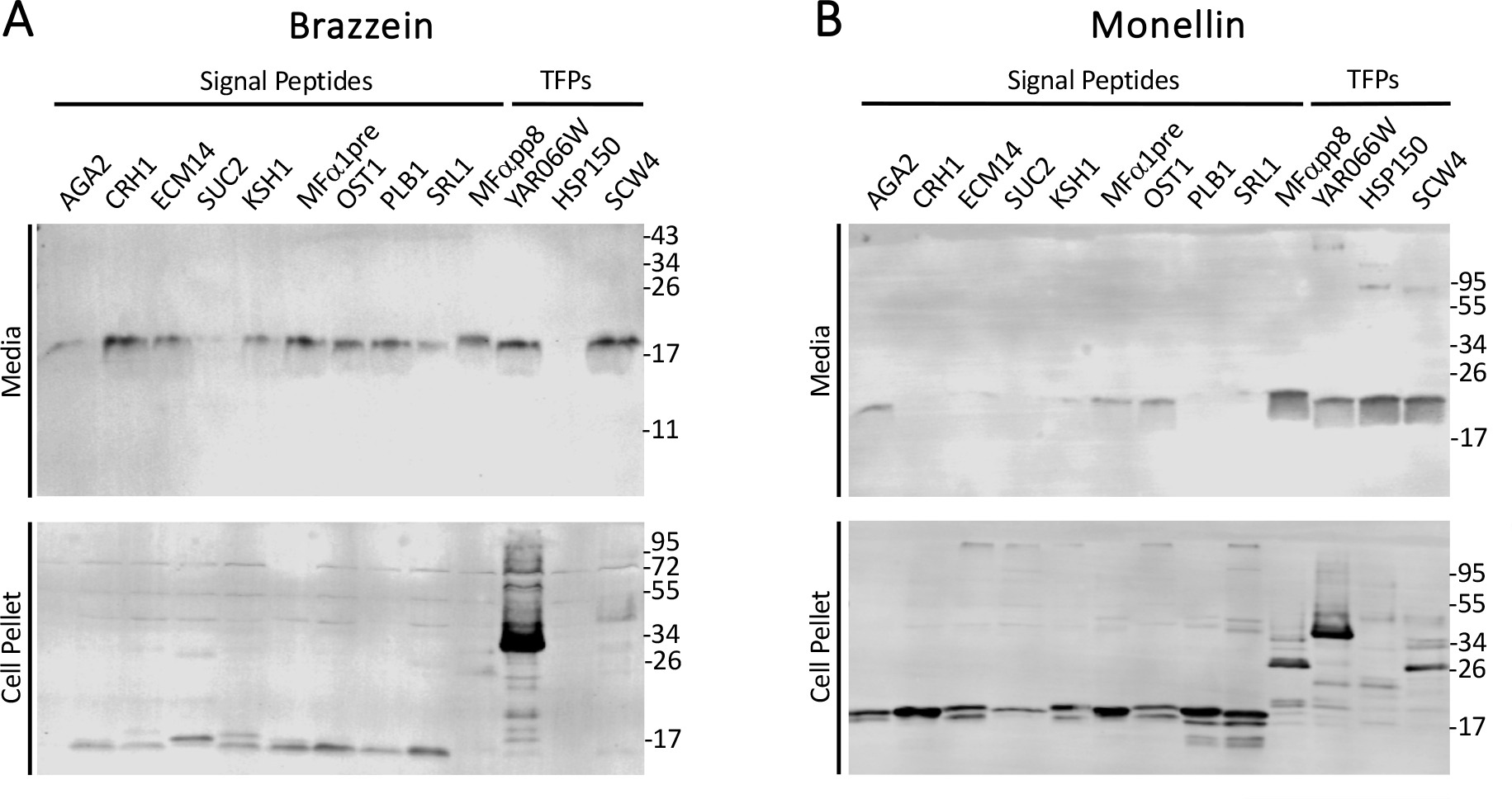
Application of the SP / TFP panel for the recombinant production of sweet proteins. Expression and secretion of recombinant sweet proteins was assessed by western blotting as before. **(A)** Most, but not all, SPs and TFPs promoted moderate levels of brazzein secretion. **(B)** Marked differences in the SPs and TFPs that gave optimal secretion of single-chain monellin were apparent. Sizes of molecular weight markers are indicated in kDa.

The second sweet protein tested was an engineered single-chain version of monellin (sc-monellin) termed MNEI *(27, 28)*. Secreted sc-monellin was detected as a broad band, slightly higher than its ∼17 kDa predicted molecular weight when expressed with pre-SPs AGA2, ECM14, KSH1, MF(α)1, OST1, and SRL1. Meanwhile, little to no secretion was observed with CRH1, SUC2 and PLB1, despite similar amounts of intracellular protein. The best levels of sc-monellin secretion were observed when it was expressed with the MFαpp8 pre-pro-SP and the TFPs, however higher molecular weight bands were observed for the TFPs suggestive of incomplete cleavage and/or heavily glycosylated forms of the protein. Intracellularly, sc-monellin was predominantly detected as well defined bands, at the correct molecular weight when expressed with the pre-SPs. Among these, a doublet was observed with AGA2, ECM14, KSH1, OST1, PLB1, and SRL1, potentially representing sc-monellin prior to and after pre-SP cleavage by signal peptidase. For MFαpp8, the TFPs and to a lesser extent for some of the pre-SPs, higher molecular weight bands were also observed intracellularly, likely reflecting differential glycosylation states of the protein.

Comparing the secretion profiles of brazzein and sc-monellin we observed variable degrees of brazzein secretion for 12 of the 13 sequences in our initial panel, while minimal if any expression or secretion seen for the TFP HSP150. For sc-monellin, on the other hand, the three TFP sequences outperformed the pre-SPs with HSP150 and MFαpp8 exhibiting the best levels of protein secretion overall. The contrasting performance of the HSP150 TFP for production of these two sweet protein underscores the difficulty of extrapolating the efficiency of a secretion promoting sequence from one protein of interest to another.

Sweet proteins have potential applications as low-calorie sweeteners. Obtaining brazzein or monellin from the native plants that produce them (*Pentadiplandra brazzeana and Dioscoreophyllum cumminsii* respectively) has not proven to be commercially viable and so there is considerable interest in their heterologous production in either established crop plants or in microorganisms amenable to industrial fermentation *(29)*. Single-chain monellin has greater thermal stability compared to the native protein consisting of two polypeptide chains. Sc-monellin was successfully secreted in *S. cerevisiae* utilising the MFα1 pre-pro-SP under the control of a GAL1 promoter *(30)*. The maximum yield achieved was 0.41 g/L, although the functionality of the secreted protein was not tested. We observed efficient secretion of monellin directed by the MFαpp8 SP, however high secretion levels were also observed for all three TFPs. Secretion levels for the TFP HSP150 were equivalent to those for MFαpp8 while showing less intracellular sc-monellin – suggesting that HSP150 might traffic sc-monellin more efficiently through the secretory pathway. There may be merit therefore in evaluating the yields of sc-monellin that could be obtained using the HSP150 or other TFPs in a larger scale fermentation system, particularly if the choice of promoter is also optimised (something which would be greatly facilitated by the promoter parts provided in the MoClo YTK). Further work would also be necessary to characterise if the resulting secreted protein is functional, especially because Liu *et al*. found that a single-chain monellin secreted using the MFα1 SP did not taste sweet likely due to glycosylation *(31)*.

Brazzein with an amino terminal Asp residue has optimal sweetness properties *(32)*. Importantly, the custom TGCT overhang used here to join SP encoding 3a’ parts to 3b’ parts encoding proteins of interests facilitates this maintenance of native amino terminal sequences following SP cleavage. Heterologous expression of brazzein has previously been achieved in transgenic plants, bacteria, *Escherichia coli* and *Lactococcus lactis*, and the yeasts, *S. cerevisiae, Pichia pastoris*, and *Kluyveromyces lactis (33)*. Brazzein was expressed intracellularly in *S. cerevisiae* and observed by western blotting at a higher than expected molecular weight of ∼35 kDa which the authors attribute to possible hyper-glycosylation *(34)*. In our hands, brazzein secreted from *S. cerevisiae* is resolved as a broad band running at, or just above, the 17 kDa. This is somewhat higher than the predicted ∼10 kDa which might suggest a degree of glycosylation, though less than that in the aforementioned report. An assessment of the glycosylation status and functionality of the secreted brazzein would therefore be of interest. Notably, while good brazzein secretion was achieved using the AGA2 and MFapp8 SPs, little or no protein was detected intracellularly. This suggests efficient targeting and passage of brazzein through the secretory pathway with these SPs – highlighting the potential value of examining both intracellular and secreted protein levels when evaluating secretion promoting sequences. In *P. pastoris*, secretion of functional brazzein was achieved utilising various SPs derived from different origins *(35)*, while in *K. lactis*, the *S. cerevisiae* MFα pre-pro- and MFα pre-SPs were used successfully to secrete functional brazzein *(36, 37)*. An existing *P. pastoris* MoClo toolkit contains a small panel of hybrid SPs that are based on the fusion of various pre-SP sequences to an engineered version of the *S. cerevisiae* MFα pre-pro-sequence *(38)*. These hybrid SPs generally did not outperform the wildtype MFα sequence, however. In addition a MoClo YTK expansion compatible with another species belonging to the *Kluyveromyces* genus *(K. marxianus)* has been described *(39)*. The broader panel of SPs and TFPs described here are compatible with these MoClo toolkits and so could potentially be applied to further optimise secretion of brazzein, or other proteins of interest, in these other biotechnologically significant yeast species.

### Adaptation of the toolkit expansion to accommodate a panel of display proteins

Aside from the production of heterologous proteins, *S. cerevisiae* is widely used for yeast surface display which has found particular application in protein engineering. Significant effort is often required to achieve efficient surface display for a given protein with the choice of anchor protein being one of the variables that may need to be optimized *(3)*. The use of modular cloning to combine our panel of SPs and TFPs with various anchor proteins has the potential to streamline this optimization process. We therefore developed a panel of five anchor proteins as type 4a parts according to MoClo YTK convention *(16)* (Fig 1C, Table 2). The panel includes the endogenous *S. cerevisiae* proteins Aga2p, Cwp2p, Sed1p and Tip1p *(40)* as well as a synthetic anchor protein called 649 stalk *(41)*. They were designed as 4a parts with a linker sequence, an HA epitope tag and a TEV protease cleavage site upstream of the anchor protein.

**Table 2.** Yeast surface display anchors used in this study.

| **Protein Name** | **Protein Description** | **#AA / MW** | **Reference** |
| --- | --- | --- | --- |
| **Aga2p** | Adhesion subunit of a-<br>agglutinin of a cells | 70 AA / 7.6 kDa | (40) |
| **Cwp2p** | Cell wall protein | 72 AA / 6.9 kDa | (40) |
| **Sed1p** | Suppression of<br>exponential defect | 320 AA / 32.7 kDa | (40) |
| **Tip1p** | Temperature shock-<br>inducible protein | 191 AA / 18.9 kDa | (40) |
| **649 Stalk** | 649 Stalk + GPI Anchor | 688 AA / 67.1 kDa | (41) |

Expression constructs for the anti-GFP nanobody were assembled for each of the anchor proteins in combination with both the AGA2 pre- and MFαpp8 SPs. Surface display was analysed by flow cytometry using an anti HA-tag antibody (Fig 4). Aga2p had to be co-expressed with its binding partner Aga1p because of low endogenous expression of Aga1p in BY4741 cells. The Aga2p, along with the Sed1p and 649 stalk anchors exhibited very efficient surface display of the nanobody for both SPs with >75% of cells positively stained. Cwp2p and Tip1p were somewhat less efficient overall but both showed notably better surface display when combined with the MFαpp8 SP than the AGA2 pre-SP. To confirm that the displayed nanobody was functional and that the linker connecting it to the anchor protein remained intact we mixed yeast cells displaying the nanobody with cell lysate from *E. coli* cells expressing GFP. Flow cytometry analysis to detect cells with bound GFP showed very similar results to those obtained using the anti HA-tag antibody for Aga2p, Sed1p and the 649 stalk. Surprisingly, for cells expressing the nanobody coupled to the Cwp2p and Tip1p anchors, labelling with GFP was much more efficient than for the anti HA-tag antibody. This may reflect greater accessibility of the smaller GFP molecules, allowing them to bind to these shorter anchor proteins. Overall, the robust binding of GFP to cells expressing the different anchor proteins demonstrates the accessibility and functionality of the displayed nanobody.

**Figure 4.**
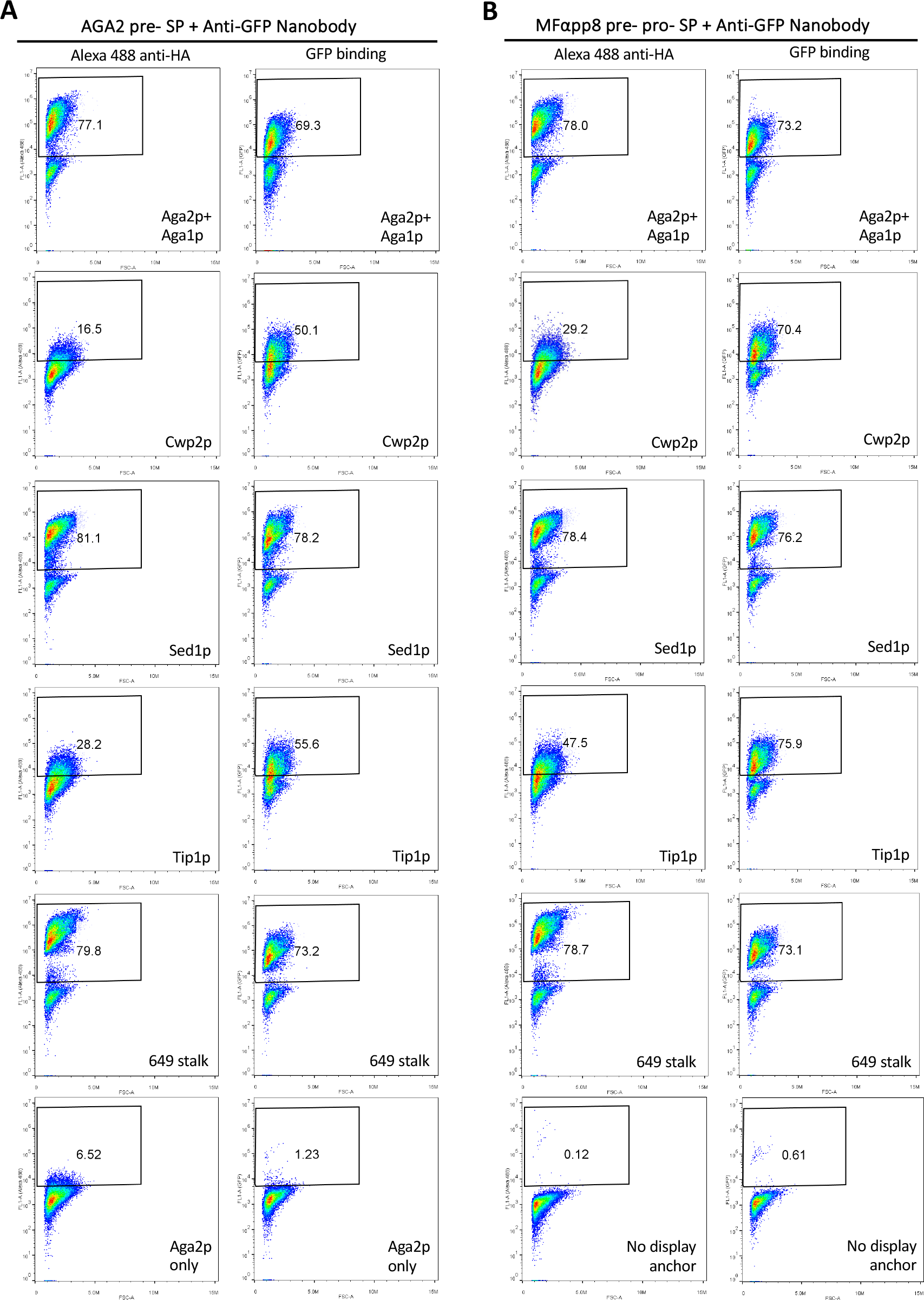
Evaluation of five anchor proteins for the surface display of an anti-GFP nanobody. The anti-GFP nanobody was expressed in yeast cells in combination with the indicated anchor proteins and either the AGA2 pre-SP **(A)** or the MFαpp8 pre-pro-SP **(B)**. Surface display was assessed by flow cytometry using an Alexa-488 fluorophore conjugated anti-HA antibody or the binding of GFP by the displayed nanobody. Untransformed BY4741 cells were used as a negative control. Plots depict FL1 signal (Alexa-488 or GFP fluorescence; Y-axis) versus forward light scattering (FSC; X-axis) with the percentage positively stained cells indicated.

We next examined the surface display of mRuby2 with each anchor protein using the AGA2 pre-SP that had worked best for mRuby2 secretion. In this case Sed1p and the 649 stalk gave efficient surface display of mRuby2, Aga2p was moderately effective, while virtually no surface display was seen with the Cwp2p and Tip1p anchors (Fig 5A). Fluorescence measurements of whole yeast cultures revealed strong mRuby fluorescence in all cases, indicating that the absence of surface display was not due to a lack of protein expression (Fig 5B).

**Figure 5.**
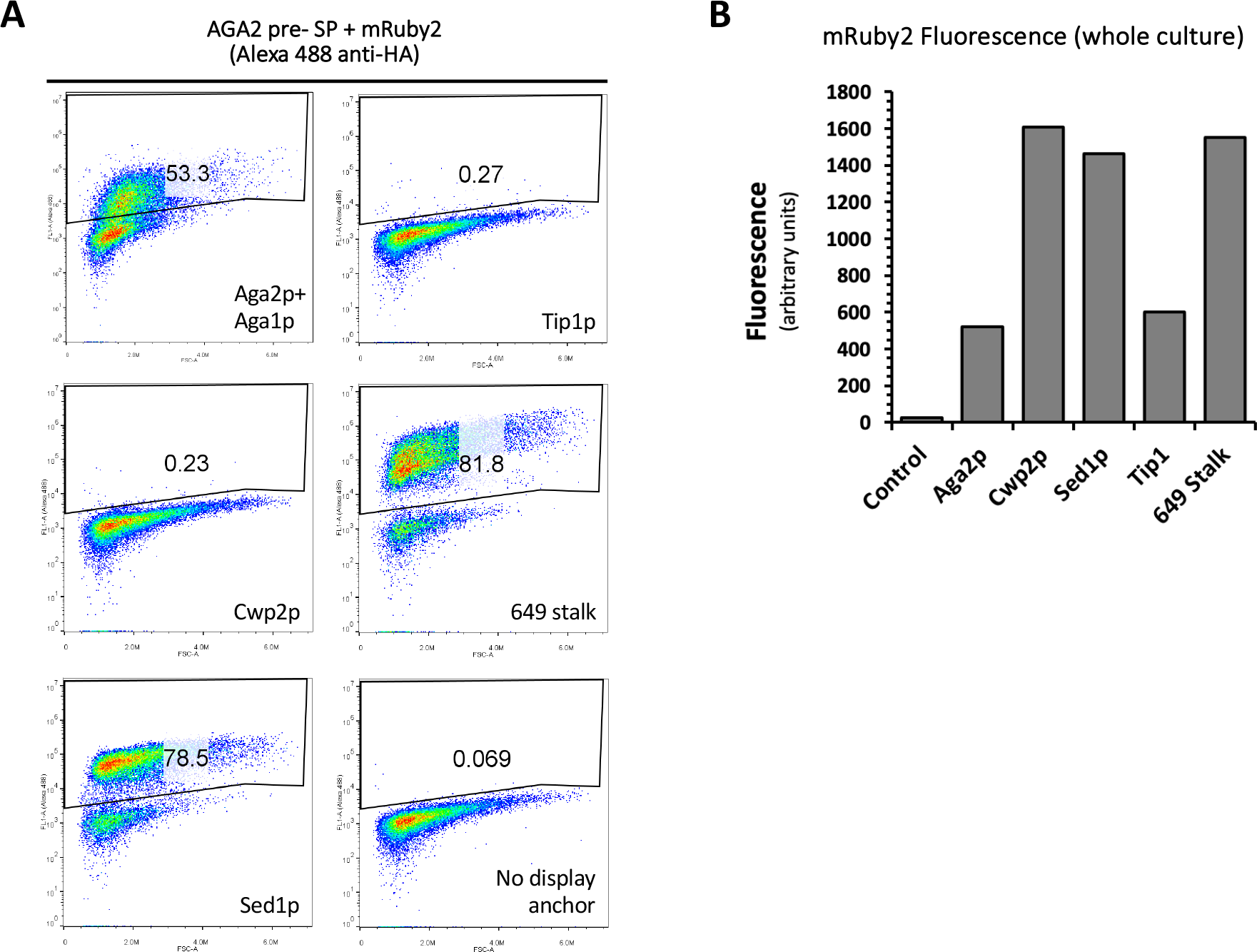
Evaluation of five anchor proteins for the surface display of mRuby2. mRuby2 was expressed in yeast cells in combination with the indicated anchor proteins and the AGA2 pre-SP. Surface display was assessed by flow cytometry using an Alexa-488 fluorophore conjugated anti-HA antibody. Untransformed BY4741 cells were used as a negative control. Plots depict FL1 signal (Alexa-488 or GFP fluorescence; Y-axis) versus forward light scattering (FSC; X-axis) with the percentage positively stained cells indicated (left panel). Measurements of mRuby2 fluorescence in whole cultures confirmed protein expression for all anchors (right panel).

Yeast surface display has long been used for protein engineering to optimize binding affinity, stability and enzymatic activity *(42)*. It has also been applied to the identification and study of protein-protein interactions, as well as the development of biosensors and whole-cell biocatalysts *(3)*. For some applications co-display of two or more proteins is required, for example when expressing multi-enzyme complexes or for co-display of an enzyme along with a protein that increases association with a substrate *(43)*. Achieving efficient display of proteins of interest, as well as controlling their expression levels and localization within the cell wall can be critical to such experiments. The panels of anchor proteins and secretion promoting sequences described here, when combined with promoters and multigene assembly capacities of the yeast modular cloning toolkit and its expansions *(16, 44)*, should accelerate the design and execution of these more sophisticated applications of yeast surface display technology.

### Summary

The MoClo YTK is a useful genetic engineering platform that permits rapid modular multipart assembly of designated DNA part types. It was originally designed to permit facile genome editing, integration, and CRISPR applications *(16)* and has since been expanded to achieve a wide range of further functionalities. Expansions include further CRISPR and genomic integration capacities, the capability to make combinatorial libraries, *(45)*, greater multiplexing capacity with more selectable markers, genomic integration sites and inducible promoters *(44)* and specific subcellular targeting of proteins *(46)*. The YSD toolkit presented here constitutes another expansion of the MoClo YTK which enables the rapid evaluation of panels of SPs and TFPs and/or anchor proteins in order to optimize the secretion and/or surface display of a protein of interest. The secretion promoting sequences and anchor proteins have been curated from the most promising candidates in the extensive literature on these topics. We suggest however, that no one universal sequence or anchor protein is likely to be suitable for every application. The prior literature, as well as the examples presented here, indicate that for many proteins of interest the optimal combination of secretion promoting sequences and/or anchor proteins must be determined empirically by experimentation and cannot be routinely predicted. For example, only three sequences from our initial panel, the SPs AGA2, OST1 and the TFP SCW4, performed well in directing secretion of all four heterologous proteins of interest that we tested. However, sequences other than these three directed the highest secretion levels of the nanobody and sc-monellin. The four heterologous proteins of interest examined have diverse structural and biochemical characteristics but are all relatively small. We expect potentially greater variability of secretion efficiency for larger, multi-domain proteins. Similarly for surface display, while the Aga2p, Sed1p and 649 stalk anchors performed consistently well, the efficiency of surface display observed for the Cwp2p and Tip1p anchors was highly dependent on the SP used and the protein of interest being displayed. Overall, these observations underline the utility of screening a range of secretion promoting sequences and/or anchors for a given protein of interest. The YSD toolkit expansion seeks to harness efficiencies of the modular cloning framework in order to standardize this trial and error process and accelerate the design, build, test cycle for synthetic biology experiments involving protein secretion or surface display in yeast.

## METHODS

### Strains and growth conditions

The *S. cerevisiae* strain used was BY4741 (MATa his3Δ1 leu2Δ0 met15Δ0 ura3Δ0). For selection, cells were grown at 30 °C in minimal synthetic media or agar plates containing 6.8 g/L yeast nitrogen base without amino acids (Sigma-Aldrich), 2 g/L dextrose (Formedium), and a complete supplement mixture of amino acids minus leucine or minus leucine and histidine (MP Biomedicals). *E. coli* DH5α cells were used for all cloning steps and were grown at 37 °C on Lysogeny Broth (LB) agar plates or in liquid LB media with shaking at 200 rpm. 35 μg/mL chloramphenicol or 50 μg/mL carbenicillin were added as appropriate.

### Chemicals, molecular biology reagents and DNA part generation

All chemicals were obtained from Merck/Sigma-Aldrich unless stated otherwise. The MoClo-YTK plasmid kit was a gift from John Dueber (Addgene kit # 1000000061). Enzymes used were from New England Biolabs unless stated otherwise. Single stranded oligonucleotides and double stranded DNAs (gBlocks) were ordered from Integrated DNA Technologies. SP, TFP, display protein and recombinant protein encoding parts were designed to be compatible with the MoClo Yeast Toolkit (YTK) *(16)*, with the exception of the overhang between 3a’ and 3b’ parts. Sequences of all parts are summarised in Supplementary File 1. Smaller parts were generated by annealing and extending complementary oligonucleotides using Klenow polymerase. Briefly, 1 μL of a 100 μM stock of each oligonucleotide were mixed with 48 μL water, heated at 95 °C for 5 minutes and then cooled by 1.5°C per minute for 1 hour. 16 μL of annealed oligonucleotide was incubated with 5 units of Klenow polymerase, 1 μL dNTPs (Thermo Fisher Scientific), and 2 μL T4 DNA Ligase Buffer at 25 °C for 15 minutes then heat inactivated at 75 °C for 20 minutes. For larger parts PCR amplification from yeast genomic DNA, plasmids or synthetic DNA fragments was performed using Phusion High Fidelity Polymerase (Thermo Fisher Scientific) according to the manufacturer’s protocol. PCR purifications, gel extractions and plasmid minipreps were performed using kits from TransGen Biotech Co. Ltd.

### MoClo DNA assembly reactions

Golden Gate Assembly reaction mixtures contained the following: ∼20 fmol of each DNA insert or plasmid, 1 μL T4 DNA ligase buffer, 0.5 μL T4 DNA ligase, 0.5 μL BsaI-HF-v2 at 20,000 units/mL or BsmBI-v2 at 10,000 units/mL, and nuclease free water to bring the total volume up to 10 μL. Assembly reactions were performed in a thermocycler. Level 0 assemblies with only one part involved 1 hour of digestion at 42 °C (BsmBI), 1 hour of ligation at 16 °C, 20 minutes of a final digestion step, and 20 minutes of heat inactivation at 80 °C. Level 0 assemblies with more than one part and level 1 assemblies involved 25-35 cycles of digestion for 2 minutes (at 37 °C for BsaI or 42 °C for BsmBI) and ligation for 5 minutes, followed by the final digestion and heat inactivation steps. A “Part 234 GFP dropout vector” pYSD099 containing ConL1, ConR2, LEU2, CEN6/ARS4 and AmpR-ColE1 parts was prepared to simplify level 1 assemblies and constitutes the backbone of most expression vectors. pYSD300, for co-expression of Aga1p, is an exception to this and has a HIS3 selectable marker gene. 2-5 uL of the assembly reaction was used for *E. coli* DH5α transformation. DNA from subsequent colonies was isolated and restriction digests were performed for verification.

### Yeast transformation

Yeast were grown at 30 °C, shaking at 200 rpm overnight in YPD media. 1 mL of pelleted cells was added to 10 μL of 10 mg/mL single stranded DNA that had been pre-incubated with 0.2–1 μg of plasmid. 500 μL LiAc-TE-PEG (100 mM Lithium Acetate, 10 mM Tris-HCL pH7.5, 1 mM EDTA, 40% w/v PEG 4000) was added to the cell pellet/DNA and incubated with occasional mixing for 15 minutes at room temperature. 50 μL DMSO was added to the tube followed by immediate heat shock at 42°C for 15 minutes with occasional mixing. Cells were pelleted, resuspended in 50 μL sterile water, and plated onto selective agar plates.

### Protein expression, western blot analysis, and fluorescence quantification

For protein expression experiments, 0.35 mL cultures of transformed cells were grown in selective synthetic media for 24 hours and then diluted to 3.5 mL in buffered YPD media (2 % bacto-tryptone, 1 % yeast extract and 2 % dextrose (Formedium) supplemented with 0.85 % MOPS free acid, and 0.1 M dipotassium phosphate, adjusted to pH 7. They were grown in 13 mL tubes with vented caps (Sarstedt) at 30°C, shaking at 200 rpm until saturated. 2 mL of the saturated culture was centrifuged at 8,000 x g for 5 minutes to pellet the cells and a 0.22 μm filter was used to ensure the secreted protein containing supernatant was cell free. 20 μL of the cell free supernatant was mixed with 6X-SDS-PAGE sample buffer. Cell pellets were lysed using the NaOH method. In brief, pellets were resuspended in 100 μL of Milli-Q water, 100 μL of 0.2 M NaOH added and incubated for 5 minutes. Samples were centrifuged at 8000 x g for 1 minute, pellets were resuspended in 50 μL 2X-SDS-PAGE sample buffer. All samples were boiled at 95°C for 5 minutes. Cell pellet samples were further diluted 1:20 in 2X-SDS-PAGE sample buffer. 10 μL of cell pellet samples and 20 μL of cell free supernatant samples was used for SDS-PAGE analysis. Western blotting was performed following transfer onto Immobilon-FL PVDF membranes (Thermo Fisher Scientific) using an anti-Histidine-tag primary antibody (Genscript A00186; 1:3000 dilution) and an IR800 conjugated secondary antibody (Li-Cor Biosciences). Images were acquired on an Odyssey imaging system and analysed using Image Studio and Empiria Studio software (Li-Cor Biosciences). Western blots shown in figures are representative of at least two independent experiments. To quantify mRuby2 fluorescence 100 μL of supernatants or whole cultures were analysed using a Tecan plate-reader with the excitation wavelength set at 559 nm and the emission wavelength set at 600 nm. In these cases, selective media was exclusively used for protein expression. Fluorescence was normalized for cell density by dividing fluorescence values by the OD_600_ values of whole cultures.

### Anti-GFP Nanobody Functionality Test

A 10 cm dish of HEK293T cells that had been transfected with the GFP expressing plasmid pEGFP-N1 (Clontech) was washed with PBS and lysed by sonication on ice in PBS supplemented with 0.2 % Triton X100, 1 mM phenylmethylsulfonyl fluoride, 0.1 mg/mL lysozyme, and 0.1 mg/mL DNAseI and then at centrifuged at 16,100 x g for 30 minutes at 4 °C to obtain the soluble cell lysate. Ni-NTA agarose beads (Neo Biotech) were prepared by washing 3 times in 1 mL PBS with centrifugation at 1000 x g, blocked in 1 % bovine serum albumin for 30 minutes at room temperature and washed once with 1 mL of equilibration buffer (20 mM Tris pH 8, 500 mM NaCl, and 5 mM Imidazole). A 50 mL saturated culture expressing the anti-GFP nanobody with the ECM14 SP was centrifuged at 4000 x g for 10 minutes to obtain the nanobody containing supernatant. 25 μL of beads were incubated for 1 hour at 4 °C with this supernatant and then washed with 3 x 1 mL PBS. The EGFP lysate was split between these beads and 25 ul of control beads that had not been incubated with the nanobody. The beads were incubated for 1 hour at 4 °C and 5 x 1 mL washes with PBS were performed. 25 μL of 200 mM imidazole was incubated with the beads for 5 minutes to elute the proteins which were mixed with 25 μL of 2X-SDS-PAGE sample buffer.

### Flow cytometry

For flow cytometry applications, 2 mL yeast cultures were grown in 13 mL tubes, in appropriate selective synthetic media at 30 °C overnight, shaking at 200 rpm. Yeast cell density was measured. Approximately 1×10^6^ cells (assuming 1 OD_600_ ≈ 1.5 x 10^7^ cells/mL) were pipetted into each microcentrifuge tube and centrifuged for 3 minutes at 3500 × g. Supernatant was removed and cells were resuspended in 100 μL of flow cytometry buffer (20 mM HEPES pH 7.5, 150 mM NaCl, 0.1% (w/v) bovine serum albumin, 5 mM maltose). Yeast were then incubated with 0.5 μg of Alexa Fluor 488-labelled HA tag antibody (Biolegend) for 15 minutes at room temperature. An unstained sample was used as a control. After incubation, cells were pelleted again, resuspended in 500 μL of buffer and analysed by flow cytometry with an Accuri C6 instrument (BD Biosciences) using the 533/30 filter and FL1 detector for green fluorescence.

### Displayed anti-GFP nanobody functionality test

25 mL of *E. coli* DH5α cells that had been transformed with the GFP Dropout Vector were grown at 37°C with shaking at 200 rpm in LB liquid media with 50 μg/mL carbenicillin. The culture was centrifuged at 3500 x g for 5 minutes at 4 °C. The pellet was resuspended in PBS supplemented with 0.2 % Triton X100, 0.1 mg/mL lysozyme, and 0.1 mg/mL DNAse and lysed by sonication on ice. The lysed cells were then centrifuged at 16,100 x g for 30 minutes at 4 °C to obtain the soluble cell lysate. The supernatant was filtered using a 0.22 μm pore size filter. For flow cytometry testing, yeast samples were prepared as described above and incubated for 15 minutes at room temperature with a 100 uL of GFP-containing supernatant. After incubation, cells were pelleted, resuspended in 500 μL of flow cytometry buffer and analysed by flow cytometry as above.

### Plasmid availability

Full details of all vectors used in this study are provided in Supplementary File 1. YTK entry vectors encoding the SP, TFP and anchor protein parts, as well as a selection of assembled constructs that can serve as controls for protein secretion or surface display, will be made available through Addgene (www.addgene.org/) as a Yeast Secretion/Display (YSD) plasmid kit.

## ABBREVIATIONS

AA: amino acid
ER: endoplasmic reticulum
FSC: forward light scattering
GFP: green fluorescent protein
GRAS: generally recognized as safe
MF: mating factor
MoClo: modular cloning
PBS: Phosphate buffered saline
POI: protein of interest
SEM: standard error of the mean
SP: signal peptides
SRP: signal recognition particle
TFP: translational fusion partners
YSD: yeast secretion/display
YTK: yeast toolkit

## AUTHOR CONTRIBUTIONS

NO’R, VJ, SO’N, AR and PY conceived and performed experiments and performed data analysis. PY conceived and supervised the project.

## CONFLICT OF INTEREST

PY is a non-renumerated director of MilisBio Ltd, who provided partial funding for the project through co-funding of an Irish Research Council Enterprise Partnership Postgraduate Scholarship to NO’R.

## ACKNOWLEDGMENTS

We thank Tadhg Crowley of the APC Flow Cytometry Platform for advice and assistance with flow cytometry. We are grateful to Dr Kate Foley, Jennifer Duane, Patricia Fowler and Dr Patrycja Zakrys for excellent technical support. We thank Dr John Morrissey, University College Cork for providing yeast strains and other reagents, and Dr John Dueber, University of California, Berkeley for the MoClo yeast toolkit obtained through Addgene. We acknowledge Joan Lenihan for her work on cloning of sweet protein sequences and Laura Cioccarelli for help with the design of the graphical abstract. This work was supported by the Irish Research Council and MilisBio Ltd through an Enterprise Partnership Postgraduate Scholarship to NO’R (Project ID: EPSPG/2020/307) and by the Science Foundation Ireland AMBER Centre for Advanced Materials and BioEngineering Research (PY and VJ; Project Number 12/RC/2278_P2).

