## Supplementary Figures for "A yeast modular cloning (MoClo) toolkit expansion for optimization of heterologous protein secretion and surface display in *Saccharomyces cerevisiae*"

**Overhang set**

CCCT,AACG,TATG,ATCC,TGGC,GCTG,TACA,GAGT,CCGA,TGCT

### Ligation frequency matrix

| | CCCT | AGGG | AACG | CGTT | TATG | CATA | ATCC | GGAT | TGGC | GCCA | GCTG | CAGC | TACA | TGTA | GAGT | ACTC | CCGA | TCGG | TGCT | AGCA |
|---|---|---|---|---|---|---|---|---|---|---|---|---|---|---|---|---|---|---|---|---|
| CCCT | 0 | 215 | 0 | 0 | 0 | 0 | 0 | 0 | 0 | 0 | 0 | 0 | 0 | 0 | 0 | 0 | 0 | 0 | 0 | 0 |
| AGGG | 215 | 0 | 0 | 0 | 0 | 0 | 0 | 0 | 0 | 0 | 0 | 0 | 0 | 0 | 0 | 0 | 0 | 0 | 0 | 0 |
| AACG | 0 | 0 | 0 | 271 | 0 | 0 | 0 | 0 | 0 | 0 | 0 | 0 | 0 | 0 | 0 | 0 | 0 | 0 | 0 | 0 |
| CGTT | 0 | 0 | 271 | 0 | 0 | 0 | 0 | 0 | 0 | 0 | 0 | 0 | 0 | 0 | 0 | 0 | 0 | 0 | 0 | 0 |
| TATG | 0 | 0 | 0 | 0 | 0 | 237 | 0 | 0 | 0 | 0 | 0 | 0 | 0 | 0 | 0 | 0 | 0 | 0 | 0 | 0 |
| CATA | 0 | 0 | 0 | 0 | 237 | 0 | 0 | 0 | 0 | 0 | 0 | 0 | 0 | 0 | 0 | 0 | 0 | 0 | 0 | 0 |
| ATCC | 0 | 0 | 0 | 0 | 0 | 0 | 0 | 284 | 0 | 0 | 0 | 0 | 0 | 0 | 0 | 0 | 0 | 0 | 0 | 0 |
| GGAT | 0 | 0 | 0 | 0 | 0 | 0 | 284 | 0 | 0 | 0 | 0 | 0 | 0 | 0 | 0 | 0 | 0 | 0 | 0 | 0 |
| TGGC | 0 | 0 | 0 | 0 | 0 | 0 | 0 | 0 | 0 | 243 | 0 | 0 | 0 | 0 | 0 | 0 | 0 | 0 | 0 | 0 |
| GCCA | 0 | 0 | 0 | 0 | 0 | 0 | 0 | 0 | 243 | 0 | 0 | 0 | 0 | 0 | 0 | 0 | 0 | 0 | 0 | 0 |
| GCTG | 0 | 0 | 0 | 0 | 0 | 0 | 0 | 0 | 0 | 0 | 0 | 173 | 0 | 0 | 0 | 0 | 0 | 0 | 0 | 0 |
| CAGC | 0 | 0 | 0 | 0 | 0 | 0 | 0 | 0 | 0 | 0 | 173 | 0 | 0 | 0 | 0 | 0 | 0 | 0 | 0 | 0 |
| TACA | 0 | 0 | 0 | 0 | 0 | 0 | 0 | 0 | 0 | 0 | 0 | 0 | 0 | 259 | 0 | 0 | 0 | 0 | 0 | 0 |
| TGTA | 0 | 0 | 0 | 0 | 0 | 0 | 0 | 0 | 0 | 0 | 0 | 0 | 259 | 0 | 0 | 0 | 0 | 0 | 0 | 0 |
| GAGT | 0 | 0 | 0 | 0 | 0 | 0 | 0 | 0 | 0 | 0 | 0 | 0 | 0 | 0 | 0 | 249 | 0 | 0 | 0 | 0 |
| ACTC | 0 | 0 | 0 | 0 | 0 | 0 | 0 | 0 | 0 | 0 | 0 | 0 | 0 | 0 | 249 | 0 | 0 | 0 | 0 | 0 |
| CCGA | 0 | 0 | 0 | 0 | 0 | 0 | 0 | 0 | 0 | 0 | 0 | 0 | 0 | 0 | 0 | 0 | 0 | 310 | 0 | 0 |
| TCGG | 0 | 0 | 0 | 0 | 0 | 0 | 0 | 0 | 0 | 0 | 0 | 0 | 0 | 0 | 0 | 0 | 310 | 2 | 0 | 0 |
| TGCT | 0 | 0 | 0 | 0 | 0 | 0 | 0 | 0 | 0 | 0 | 0 | 0 | 0 | 0 | 0 | 0 | 0 | 0 | 1 | 239 |
| AGCA | 0 | 0 | 0 | 0 | 0 | 0 | 0 | 0 | 0 | 0 | 0 | 0 | 0 | 0 | 0 | 0 | 0 | 0 | 239 | 0 |

**Legend**

- good Watson-Crick pair
- poor Watson-Crick pair
- high-count mismatch
- modest mismatch
- trace mismatch

**Supplementary Figure 1. Analysis of the TGCT overhang when used as part of YTK toolkit using the NEBridge GetSet tool from New England Biolabs** The overhangs from the yeast modular cloning toolkit were analysed with TGCT replacing the standard Part 3a/3b overhang. Minimal mismatches are observed and Golden Gate Assembly is predicted to yield 99% of correctly-ligated products using this set of overhangs.

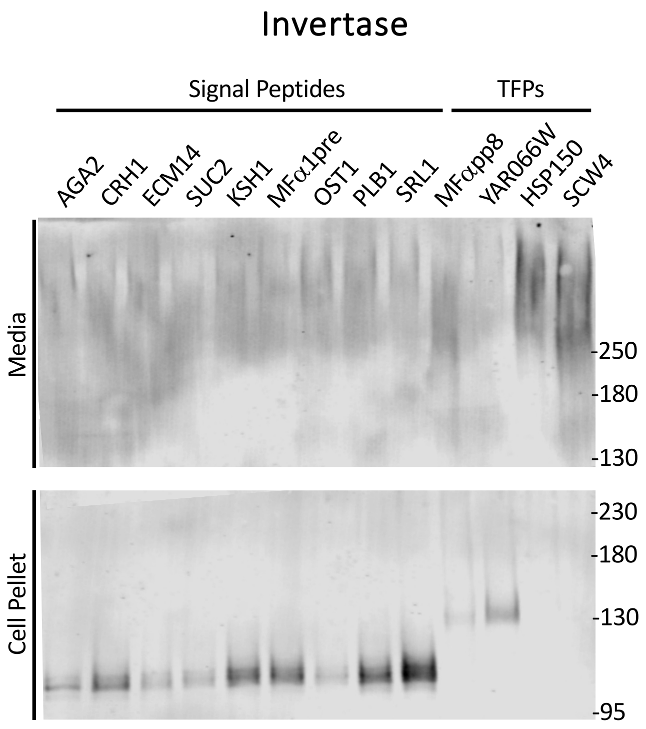


**Supplementary Figure 2. Evaluation of a panel of thirteen secretion promoting sequences for the expression of yeast invertase.** Recombinant proteins were targeted for secretion using the indicated SPs and TFPs. Secreted and intracellular proteins were detected from the media and cell pellet respectively by western blotting for a carboxyl-terminal 6xHis tag. Secreted invertase was detected for all SP and TFP sequences as a very high molecular weight smear – likely due to extensive glycosylation.
